# Gigantic Genomes Can Provide Empirical Tests of TE Dynamics Models — An Example from Amphibians

**DOI:** 10.1101/2020.08.19.257527

**Authors:** Jie Wang, Michael W. Itgen, Huiju Wang, Yuzhou Gong, Jianping Jiang, Jiatang Li, Cheng Sun, Stanley K. Sessions, Rachel Lockridge Mueller

## Abstract

Transposable elements (TEs) are a major determinant of eukaryotic genome size. The collective properties of a genomic TE community reveal the history of TE/host evolutionary dynamics and impact present-day host structure and function, from genome to organism levels. In rare cases, TE community/genome size has greatly expanded in animals, associated with increased cell size and altered anatomy and physiology. We characterize the TE landscape of the genome and transcriptome in an amphibian with a giant genome — the caecilian *Ichthyophis bannanicus*, which we show has a genome size of 12.2 Gb. Amphibians are an important model system because the clade includes independent cases of genomic gigantism. The *I. bannanicus* genome differs compositionally from other giant amphibian genomes, but shares a low rate of ectopic-recombination-mediated deletion. We examine TE activity using expression and divergence plots; TEs account for 15% of somatic transcription, and most superfamilies appear active. We quantify TE diversity in the caecilian, as well as other vertebrates with a range of genome sizes, using diversity indices commonly applied in community ecology. We synthesize previous models integrating TE abundance, diversity, and activity, and we test whether the caecilian meets model predictions for genomes with high TE abundance. We propose thorough, consistent characterization of TEs to strengthen future comparative analyses. Such analyses will ultimately be required to reveal whether the divergent TE assemblages found across convergent gigantic genomes reflect fundamental shared features of TE/host genome evolutionary dynamics.

## Introduction

Transposable elements (TEs) are segments of DNA that move within genomes [1]. Because their movement is often associated with an increase in copy number, these elements constitute a substantial — but variable — fraction of eukaryotic genomes, e.g. 2.7% in pufferfish (*Takifugu rubripes*) [2] and 85% in maize (*Zea mays*) [3]. TEs were discovered by Barbara McClintock in the late 1940s, demonstrating that genomes are far more dynamic entities than previously thought [4].

Although they share the characteristic of intra-genomic mobility, TEs are highly diverse sequences. TE classification has been updated over the years to reflect new discoveries [5]. Several classification systems have been proposed that establish nested groups according to transposition mechanism, structure, sequence similarity, and, to some degree, shared evolutionary history [6-10]. These classification systems, in turn, have allowed the community of genome biologists to annotate TEs in the genomes of diverse species, identifying differences in overall TE composition, TE activity, TE turnover dynamics, and TE domestication across the tree of life [11-13].

Overall TE content is the main predictor of haploid genome size, which shapes a variety of traits including the sizes of nuclei and cells, the rates of development and basal metabolism, and the structural complexity of organs [14-21]. The evolutionary forces that shape TE load include: mutation (specifically the insertion of new TE sequences by transposition and their removal by deletion), selection (which likely targets individual TE loci as well as the pathways that control TE transposition and deletion) [22], and genetic drift (which determines how efficiently purifying selection purges deleterious TE sequences) [23]. How these forces interact to generate genome size diversity across the tree of life remains incompletely understood. Groups of related species that vary in TE load and genome size provide critical model systems for studying this fundamental question [24].

Across animals, genomic gigantism is rare. Within vertebrates, it is best understood in a group related to the caecilians — the salamanders (order Caudata), a clade of ∼700 extant species of amphibians (almost exclusively diploid) with haploid genome sizes that range from 14 Gb to 120 Gb [25]. Fossil cell size data demonstrate that salamander genome sizes have been large for ∼160 million years [26]. Comparative genomic analyses demonstrate that salamander genomes have high levels of TEs — particularly LTR retrotransposons — and that these high levels reflect low rates of DNA loss in dead-on-arrival non-LTR retrotransposons, low rates of ectopically-mediated LTR retrotransposon deletion, and intact piRNA-mediated TE silencing machinery, albeit with fewer TE-targeting piRNAs than are found in animals with smaller genomes [27-33]. Phylogenetic comparative analyses demonstrate that salamanders’ enormous genomes result from an abrupt change in evolutionary dynamics at the base of the clade, implying a discrete shift in the balance among the evolutionary forces shaping TE accumulation [34, 35].

There are three living clades of amphibians: caecilians (order Gymnophiona), salamanders (order Caudata), and frogs and toads (order Anura); Caudata and Anura are sister taxa, and Gymnophiona is the sister taxon to Caudata + Anura. Frogs and toads are a well-studied group of 7175 species. Of the 278 species (in 78 genera) for which genome size estimates exist, a handful of species in three different genera have genomes that reach or exceed 10 Gb [25], providing independent examples of genomic expansion. Genomic data examined to date show diverse TE landscapes across species [36-41], but no sequence data exist (to our knowledge) for those with the largest genomes. Caecilians are a relatively understudied group of 214 extant species, all of which are limbless, serpentine, burrowing or aquatic animals with reduced eyes, ringed bodies, and strong, heavily ossified skulls. Genome size estimates exist for roughly 20 species and range from 2.8 Gb to 13.7 Gb [25, 35]. These data show yet another independent example of genomic expansion within amphibians, suggesting a clade-wide propensity towards TE accumulation relative to other vertebrates that is most extreme in salamanders. Genomic data for caecilians are sparse, but growing based on successes of the G10K consortium and others [42, 43]. Published data are lacking for species with the largest genomes. This lack of amphibian data underlies a major gap in our knowledge of vertebrate genome evolution [13]. More generally, a lack of detailed information on TE biology in large and repetitive genomes, reflecting persistent assembly and annotation challenges, underlies a major gap in genome biology as a whole.

In this study, we present an analysis of TE biology in a caecilian with a large genome — *Ichthyophis bannanicus* (Gymnophiona: Ichthyophiidae), which we show has a genome size of 12.2 Gb. We compare the caecilian to other vertebrates with diverse genome sizes, demonstrating how the TE community in a large genome can be used to evaluate existing models of TE dynamics. *I. bannanicus* is a relatively small species (adult sizes 30-41 cm) with an aquatic larval stage and terrestrial/fossorial adult stage. Its distribution includes China and northern Vietnam, and it is an IUCN species of Least Concern. We analyzed both genomic shotgun sequence data and RNA-Seq data from diverse tissues to answer the following specific questions: (1) what abundance and diversity of TEs make up the bulk of the large *I. bannanicus* genome? (2) what are the amplification and deletion dynamics of TEs in the genome? (3) what contribution does the large genomic TE load make to the somatic transcriptome? (4) do the patterns of genomic TE composition and overall TE expression fit the predictions of models of TE dynamics in large genomes? We show that up to 68% of the *I. bannanicus* genome is composed of TEs, with another 9% identified as repetitive sequences not classifiable as known TEs. The two most abundant TE superfamilies — DIRS/DIRS and LINE/Jockey — account for ∼50% of the genome. Unlike salamander genomes, the *I. bannanicus* genome has relatively few LTR retrotransposons, demonstrating that repeated instances of extreme TE accumulation in amphibians do not reflect failure to control a specific type of TE. We show that rates of ectopic-recombination-mediated deletion are low relative to vertebrates with more typical genome sizes, and that TE expression is high. We quantify and compare TE diversity in *I. bannanicus* and 11 other vertebrates using indices common in community ecology. We demonstrate that comparative analyses of TE diversity can be a powerful tool for evaluating models of TE dynamics, and we show that it could be even more powerful if researchers adopt a uniform approach to TE diversity analysis. We propose such an approach to move the field forward. Taken together, our results demonstrate that computationally feasible analyses of large genomes can reveal the genomic characteristics favoring expanded TE communities, as well as the resulting impact of high TE load on the transcriptome. Such analyses targeting phylogenetically diverse organisms can yield fundamental insights into the complex ways in which TEs drive genome biology.

## Results

### The *I. bannanicus* genome is 12.2 Gb and contains most known TE superfamilies

The haploid genome size of *I. bannanicus* was estimated to be 12.2 Gb based on analyses of Feulgen-stained erythrocytes following established methods [14]. This estimate is similar to the other published estimate from the same genus (*I. glutinosus*, 11.5 Gb) [35]. We used the PiRATE pipeline [44], designed to mine and classify repeats from low-coverage genomic shotgun data in taxa — such as caecilians — that lack genomic resources. The pipeline yielded 59,825 contigs (**Table 1**). RepeatMasker mined the majority of the repeats (37,123 out of 59,825, or 62.1%). dnaPipeTE was the second most effective tool, mining 19,160 repeats (32.0%), followed by RepeatScout (3.0%) and TE-HMMER (2.7%). In this pipeline, TE-denovo, LTR-harvest, Helsearch, SINE-finder, and MITE hunter found few additional repeats, and we found no additional repeats using MGEScan. Clustering with CD-HIT-est with a 95% sequence identity cutoff yielded 51,862 contigs, and clustering at 80% yielded 23,092 contigs.

**Table 1.** Repeat contigs identified by different methods/software of the PiRATE pipeline.

| **TE-mining method** | **Software** | **Repeats clustered at 100% identify** |
| --- | --- | --- |
| Similarity-based | RepeatMasker | 37,123 (62.1%) |
|  | TE-HMMER | 1,585 (2.7%) |
| Structure-based | HelSearch | 17 (0.0%) |
|  | LTR harvest | 23 (0.0%) |
|  | MGEScan | 0 |
|  | MITE hunter | 12 (0.0%) |
|  | SINE finder | 17 (0.0%) |
| Repetitiveness-based | TEdenovo | 102 (0.2%) |
|  | RepeatScout | 1,786 (3.0%) |
| Repeat-building-based | dnaPipeTE | 19,160 (32.0%) |
| In total |  | 59,825 (100%) |

Repeat contigs were annotated as transposable elements to the levels of order and superfamily in Wicker’s hierarchical classification system [7], modified to include several recently discovered TE superfamilies, using PASTEC [45]. Of the 59,825 identified repeat contigs, 50,471 (84.4%) were classified as known TEs (**Table 2**). Transposable elements representing eight of the nine orders proposed in Wicker’s system are present in the *I. bannanicus* genome; only Crypton was not identified by our pipeline (although we note that 192 chimeric contigs were filtered out that included a Crypton annotation, and nine transcriptome contigs were annotated as Crypton). Within these eight orders, our analyses identified 25 TE superfamilies, each represented by between two and 26,507 annotated contigs. Non-autonomous TRIM, LARD, and MITE elements are also present in the *I. bannanicus* genome, represented by 229, 28, and 146 contigs, respectively, and an additional 277 contigs were only annotated to the level of order or class (i.e. unknown LINE, SINE, and TIR or unknown Class I) (Table 2).

**Table 2.**
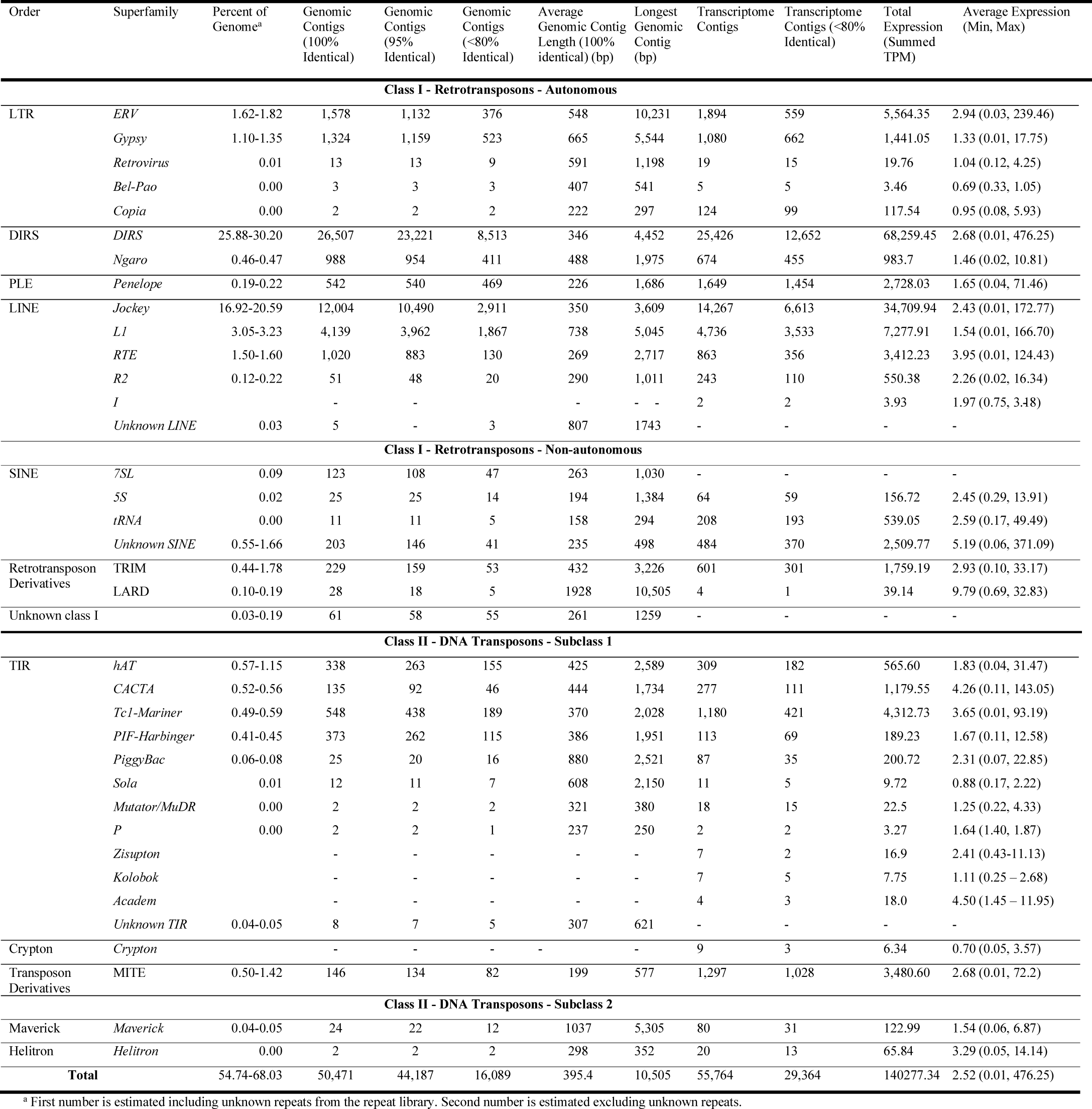
Classification of repeat contigs and summary of repeats detected in the genome and somatic transcriptome.

### 78% of the *I. bannanicus* genome is repetitive, dominated by DIRS elements

To calculate the percentage of the caecilian genome composed of different TEs, the shotgun reads were masked with RepeatMasker v-4.0.7 using our caecilian-derived repeat library. We then repeated the RepeatMasker analysis excluding the unknown repeats and compared the two sets of results as a rough approximation of the number of unknown repeat contigs that were, in fact, TE-derived sequences that were divergent, fragmented, or otherwise unidentifiable by our pipeline. 68.2% of these sequences (measured as bp) were masked as repetitive when the repeat library included only the 50,471 contigs classified as TEs and the 29 contigs annotated as putative multi-copy host genes; 66.1% were identifiable to the superfamily level of TEs (Table 2), an additional 1.94% were identifiable only to the class or order level, and 0.17% were multi-copy host genes. When the analysis was performed including the 9325 unknown-repeat contigs, along with the classified TEs and putative multi-copy host genes, 77.6% of the data were masked as repetitive overall, suggesting that the unknown repeats comprise 9.4% of the genome. However, the percentage of the genome identified as known TEs decreased from 68.0% to 54.7% with the inclusion of unknown repeats, demonstrating that many reads were sufficiently similar to known TEs to be masked by them when unknown repeat contigs were not available as a best-match option. This result suggests that at least some of the unknown repeats are TE-derived sequences.

Class I TEs (retrotransposons) make up 52.09-63.68% (unknown repeats included or excluded in the repeat library, respectively) of the *I. bannanicus* genome; they are almost 20 times more abundant than Class II TEs (DNA transposons; 2.63–4.36%). DIRS/DIRS is the most abundant superfamily (25.88-30.20% of the genome), followed by LINE/Jockey (16.92-20.59%), LINE/L1 (3.05-3.23%), LTR/ERV (1.62-1.82%), LINE/RTE (1.50-1.60%), and LTR/Gypsy (1.10-1.35%); all are retrotransposons (Table 2). TIR/hAT (0.57-1.15%), TIR/CACTA (0.52-0.56%), and TIR/Tc1-Mariner (0.49-0.59%) are the most abundant superfamilies of DNA transposons (Table 2). These proportions differ from those found in the gigantic genomes of salamanders, where LTR/Gypsy elements dominate (7 – 20% of the genome, depending on species), DIRS/DIRS elements never exceed 7% of the genome, and LINE/Jockey elements never exceed 0.03% of the genome [30, 32]. Here and throughout the paper, we are interpreting our results based on the assumption that the genomic shotgun data are a random representation of the whole genome; Illumina reads should sample the genome in a random and independent manner, despite some stochastic sampling error.

### The *I. bannanicus* genome shows low diversity index values when measured at the TE superfamily level

Diversity indices are mathematical measures of diversity within a community. In ecology, they are widely used to summarize species diversity within an ecological community, although they are also used in other fields (e.g. economics). Diversity indices take into account species richness (the total number of species present) and evenness (based on the proportional abundance of each species) [46]. Within genome biology, richness can summarize the total number of TE types (e.g. TE superfamilies) and evenness can summarize the proportion of the genome occupied by each TE type [47-50]. We calculated two commonly used diversity indices — the Shannon Index and the Gini-Simpson Index [51, 52] — on the caecilian TE community, as well as the TE communities from 11 other vertebrates spanning a range of genome sizes and types of datasets. Genome sizes ranged from 0.4 Gb (the pufferfish *Takifugu rubripes*) to 55 Gb (the hellbender salamander *Cryptobranchus alleganiensis*). Datasets ranged from full genome assemblies to low-coverage genome skims. The Shannon Index quantifies the uncertainty in identity of an individual drawn at random from a community. The Gini-Simpson index quantifies the probability that two individuals drawn at random from a community are different types, and it gives more weight to dominant (i.e. most abundant) species. Results are summarized in **Table 3**. The Shannon Index ranges from 0.9 (chicken, the least diverse) to 2.41 (green anole lizard, the most diverse). The Gini-Simpson Index ranges from 0.5 (chicken, the least diverse) to 1 (pufferfish, the most diverse). By both indices, the caecilian has the second-least diverse genome of the 11 total genomes compared. There is no overall correlation between genome size and TE diversity using either index.

**Table 3.** Diversity indices summarizing the TE communities from 11 vertebrate genomes.

| Species | Common Name | Genome Size (Gb) | Shannon Index | Gini-Simpson Index | Dataset |
| --- | --- | --- | --- | --- | --- |
| *Takifugu rubripes* | Pufferfish | 0.4 | 2.10 | 1.00 | Full genome assembly |
| *Gallus gallus* | Chicken | 1.3 | 0.90 | 0.50 | Full genome assembly |
| *Xenopus tropicalis* | Western clawed frog | 1.7 | 2.24 | 0.90 | Full genome assembly |
| *Anolis carolinensis* | Green anole lizard | 2.2 | 2.41 | 0.91 | Full genome assembly |
| *Homo sapiens* | Human | 3.1 | 1.69 | 0.79 | Full genome assembly |
| *Ichthyophis bannanicus* | Banna caecilian | 12.2 | 1.45 | 0.67 | ~0.1X genome skimming |
| *Desmognathus ochrophaeus* | Allegheny Mountain dusky salamander | 15 | 1.61 | 0.71 | ~0.01X genome skimming |
| *Batrachoseps nigriventris* | Black-bellied slender salamander | 25 | 2.18 | 0.86 | ~0.01X genome skimming |
| *Ambystoma mexicanum* | Mexican axolotl salamander | 32 | 2.26 | 0.89 | Assembly |
| *Aneides flavipunctatus* | Speckled black salamander | 44 | 1.96 | 0.78 | ~0.01X genome skimming |
| *Cryptobranchus alleganiensis* | Hellbender salamander | 55 | 2.02 | 0.84 | ~0.01X genome skimming |

### Most TE superfamilies are active in the *I. bannanicus* genome

For each of the 19 TE superfamilies accounting for ≥ 0.005% of the genome, the overall amplification history was summarized by plotting the genetic distances between individual reads (representing TE loci) and the corresponding ancestral TE sequences as a histogram with bins of 1%. Of these 19 TE superfamilies, 17 of the resulting distributions showed characteristics of ongoing or recent activity (i.e. presence of TE sequences <1% diverged from the ancestral sequence and a unimodal right-skewed, J-shaped, or monotonically decreasing distribution) (**Figure 1**). Six of these showed essentially unimodal, right-skewed distributions: LTR/ERV, DIRS/DIRS, LINE/Jockey, LINE/RTE, TIR/PiggyBac, and TIR/Sola. An additional three showed essentially unimodal, right-skewed distributions with a spike in sequences <1% diverged from the ancestral sequence: SINE/7SL, TIR/hAT, and TIR/Tc1/mariner. A single superfamily — PLE/Penelope — showed a left-skewed J-shaped distribution. These ten distributions suggest TE superfamilies that continue to be active today, but whose accumulation peaked at some point in the past. In contrast, six TE superfamilies showed essentially monotonically decreasing distributions with a maximum at <1% diverged from the ancestral sequence: LTR/Gypsy, DIRS/Ngaro, LINE/L1, TIR/CACTA, TIR/PIF-Harbinger, and Maverick. SINE/5S has a bimodal distribution with a maximum at <1% diverged from the ancestral sequence. These seven distributions suggest TE superfamilies that continue to be active today at their highest-ever rates of accumulation. Two superfamilies — LTR/Retrovirus and LINE/R2 — appear largely inactive, showing unimodal distributions with few sequences <1% diverged from the ancestral. For almost all superfamilies, multiple contigs that were <80% identical in sequence to one another were assembled (range 1 – 8513), suggesting the presence of many families within each superfamily.

**Figure 1.**
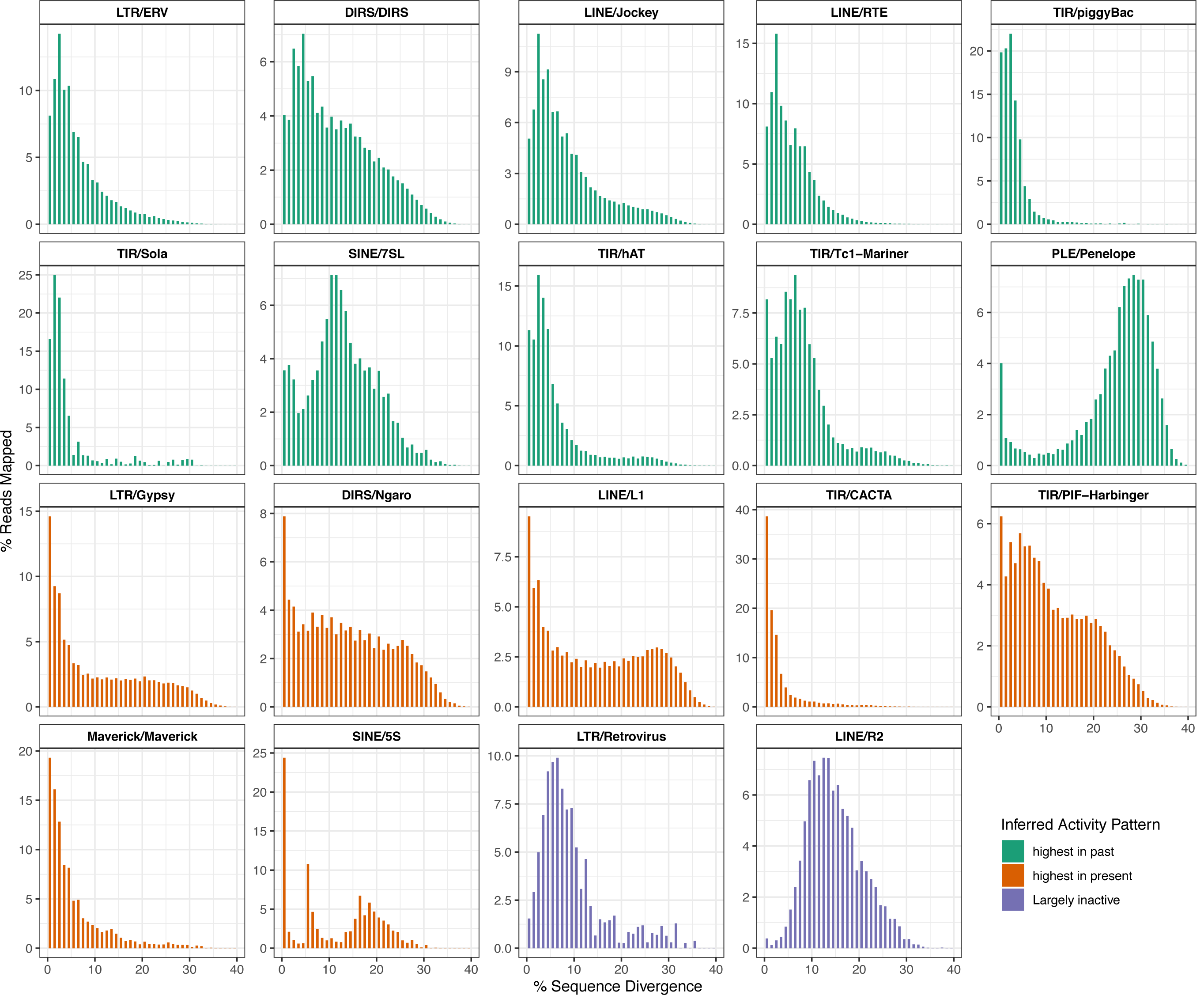
Amplification plots for transposable element, TE, superfamilies. The majority (17/19) suggest current superfamily activity. Note that the y-axes differ in scale.

### Ectopic-recombination-mediated deletion levels are lower in *I. bannanicus* than in vertebrates with smaller genomes

Ectopic recombination, also known as non-allelic homologous recombination, occurs between two DNA regions that are similar in sequence, but do not occupy the same locus. Ectopic recombination among LTR retrotransposon sequences can produce deletions that leave behind solo LTRs, which are single terminal repeat sequences that lack the corresponding internal sequence and matching terminal repeat sequence. Accordingly, the ratio of LTR sequences to internal retrotransposon sequences can be used to estimate levels of ectopic-recombination-mediated deletion. Larger genomes like *I. bannanicus* are predicted to have lower levels of deletion [33].

Two superfamilies were selected for ectopic recombination analysis: DIRS/DIRS, which accounts for over a quarter of the caecilian genome, and LTR/Gypsy, which is one of the two most abundant LTR superfamilies in the caecilian genome at 1.4%, but which dominates other gigantic amphibian genomes [53]. Mean estimates of the total terminal sequence to internal sequence ratio (TT:I) across the 9 DIRS/DIRS contigs range from 1.2:1 to 0.7:1, depending on the minimum alignment length for reads (**Figure 2**). Mean TT:I estimates across the 17 LTR/Gypsy contigs range from 1.3:1 to 1.2:1. Values of 1:1 are expected in the absence of ectopic-recombination-mediated deletion. The higher sensitivity of DIRS/DIRS than LTR/Gypsy to the minimum alignment length parameter value likely reflects the shorter length of the terminal sequence (150 bp vs. 744 bp in DIRS/DIRS and LTR/Gypsy, respectively); the 0.7:1 TT:I value for DIRS/DIRS is likely an underestimate. Variation in the TT:I ratio among contigs in each superfamily was similar (Figure 2) and lower than the ranges reported in vertebrates with more typically sized (i.e. smaller) genomes [33].

**Figure 2.**
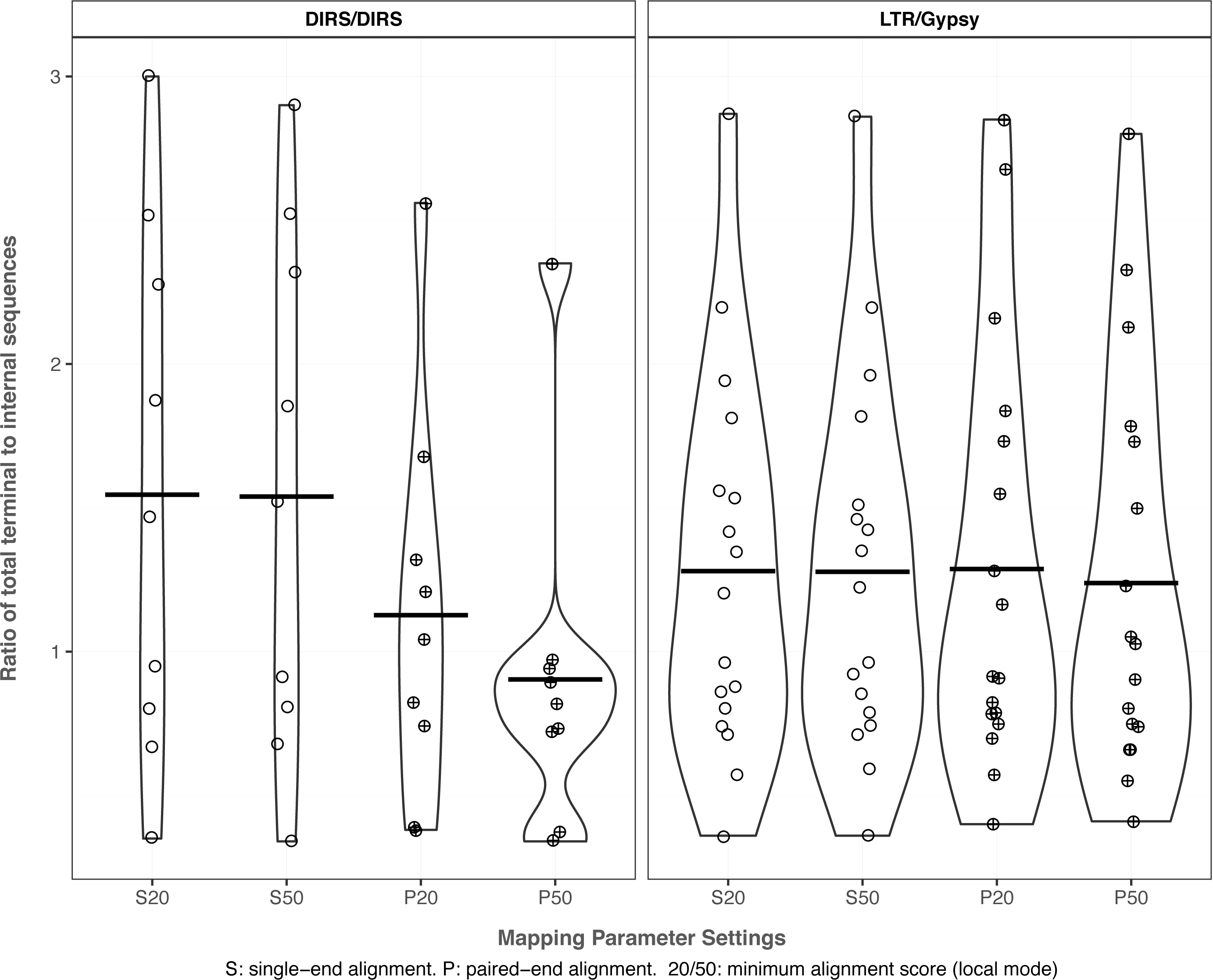
Ratio of total terminal sequence to internal sequence for two TE superfamilies. A ratio of 1:1 is expected in the absence of ectopic-recombination-mediated deletion. S: single−end alignment. P: paired−end alignment. 20/50: minimum alignment score (local mode).

For both superfamilies, ectopic-recombination-mediated deletion levels in the caecilian (TT:I ratio ∼1.2:1) are similar to the low levels estimated from four gigantic salamander genomes (TT:I ratios 0.55:1 to 1.25:1 for *Aneides flavipunctatus, Batrachoseps nigriventris, Bolitoglossa occidentalis*, and *Bolitoglossa rostrata*) and below the levels estimated from vertebrates with more typically sized (i.e. smaller) genomes (TT:I ratios 1.7:1 to 3.35:1 for *Anolis carolinensis, Danio rerio, Gallus gallus, Homo sapiens*, and *Xenopus tropicalis* [33]. TT:I ratios measured for LTR/Gypsy in two salamander species (*A. flavipunctatus* and *B. nigriventris*) are 0.9:1 and 1.25:1, encompassing the value for *I. bannanicus* LTR/Gypsy.

If deletion levels were the same between the two superfamilies in the *I. bannanicus* genome, the DIRS/DIRS TT:I ratio would be expected to be lower than the LTR/Gypsy TT:I ratio because of the structure of DIRS/DIRS; it has inverted terminal repeats and internal complementary regions [54, 55] that are expected to produce incomplete deletion of the internal sequence following ectopic recombination. The higher TT:I ratio actually estimated in DIRS/DIRS may reflect the greater abundance of this superfamily, which increases the number of potential off-targets for recombination, offsetting both the incomplete deletion of the internal sequence as well as the shorter terminal sequences in DIRS that would predict lower levels of deletion [56].

### Autonomous and non-autonomous TEs are transcribed in *I. bannanicus*

To annotate transcriptome contigs containing autonomous TEs (i.e. those with open reading frames encoding the proteins necessary for transposition), BLASTx was used against the Transposable Element Protein Database (RepeatPeps.lib, http://www.repeatmasker.org/). To annotate contigs containing non-autonomous TEs that lack identifiable open reading frames, RepeatMasker was used with our caecilian-derived genomic repeat library of non-autonomous TEs. To identify contigs that contained an endogenous caecilian gene, the Trinotate annotation suite was used [57]. 38,584 contigs were annotated as endogenous (i.e. non-TE-derived) caecilian genes. 53,106 contigs were annotated as autonomous TEs using BLASTx against the Transposable Element Protein Database (RepeatPeps.lib). An additional 2658 contigs were annotated as non-autonomous TEs using the caecilian TRIM-, LARD-, SINE- and MITE-annotated genomic contigs. 1445 contigs were annotated as both autonomous TEs and endogenous caecilian genes, and an additional 342 were annotated as both non-autonomous TEs and endogenous caecilian genes (**Table 4**).

**Table 4.** Overall summary of transcriptome annotation (contigs with TPM ≥ 0.01)

|  | Contigs<br>(Percentage of<br>total contigs) | Summed TPM<br>(Percentage of<br>total expression) | Maximum<br>TPM | Minimum<br>TPM | Average<br>TPM | Mean Contig<br>Length (bp) |
| --- | --- | --- | --- | --- | --- | --- |
| Endogenous genes | 38,584 (13.3%) | 295,759 (29.6%) | 10,112.76 | 0.01 | 7.7 | 2,086 |
| Autonomous TEs | 53,106 (18.4%) | 131,793 (13.2%) | 476.25 | 0.01 | 2.5 | 570 |
| Non-Autonomous TEs | 2,658 (0.9%) | 8,484 (0.9%) | 371.09 | 0.01 | 3.2 | 785 |
| Gene/Autonomous TE | 1,445 (0.5%) | 4,859 (0.5%) | 383.94 | 0.01 | 3.7 | 2,161 |
| Gene/Non-autonomous TE | 342 (0.1%) | 1,584 (0.2%) | 198.8 | 0.01 | 4.6 | 2,800 |
| Unannotated | 193,245 (66.8%) | 555,776 (55.6%) | 8,224.66 | 0.01 | 2.9 | 537 |
| Total | 289,380 (100%) | 1,000,006 (100%) | 10,112.76 | 0.01 | 3.5 | 763 |

Of the 20 most highly expressed putative “TE/gene” contigs, ten were confirmed to have annotations for both a TE and a gene with non-overlapping ORFs. Of these, the TE was upstream of the gene in eight cases and downstream in two cases. Six of the upstream TEs were autonomous and thus contained ORFs; four of these were encoded on the same strand as the gene (two in-frame, two not in-frame) and two were encoded on the opposite strand. One of the two downstream TEs was autonomous, and it was encoded on the opposite strand from the gene. Although requiring further validation, these results suggest that at least some gene/TE pairs are co-transcribed, a way in which TE insertions can regulate gene expression [58]. One contig had overlapping annotations of a gene and a TE, a pattern that could reflect either convergence in sequence or exaptation of a TE [59].

### TE expression correlates with genomic abundance in *I. bannanicus*

Among the transcriptome contigs with TPM (transcripts per million) ≥ 0.01, autonomous TEs account for 18.4% of the total transcriptome contigs and 13.2% of the overall somatic transcriptome (TPM = 131,793) (Table 4, **Figure 3**). Non-autonomous TEs account for 0.9% of the total transcriptome contigs and 0.8% of the somatic transcriptome (summed TPM = 8484). Contigs annotated both as TEs and endogenous caecilian genes account for 0.6% of annotated transcriptome contigs and 0.6% of the somatic transcriptome (summed TPM = 6443). Endogenous (non-TE-derived) caecilian genes account for 13.3% of the total transcriptome contigs and 29.6% of the somatic transcriptome (summed TPM = 295,759). Unannotated contigs account for 66.8% of the total transcriptome contigs and 55.6% of the somatic transcriptome (summed TPM of unannotated contigs = 555,776). Five superfamilies (*I, Zisupton, Kolobok, Academ*, and *Crypton*) were detected at low expression levels in the transcriptome, but were not initially detected in the genomic data; mapping the genomic reads to these transcriptome contigs with Bowtie2 identified ≤ 3 reads per superfamily, indicating their extremely low frequency in the genome. In contrast, only one superfamily (*7SL*) was detected in the genomic data but not the transcriptome data.

**Figure 3.**
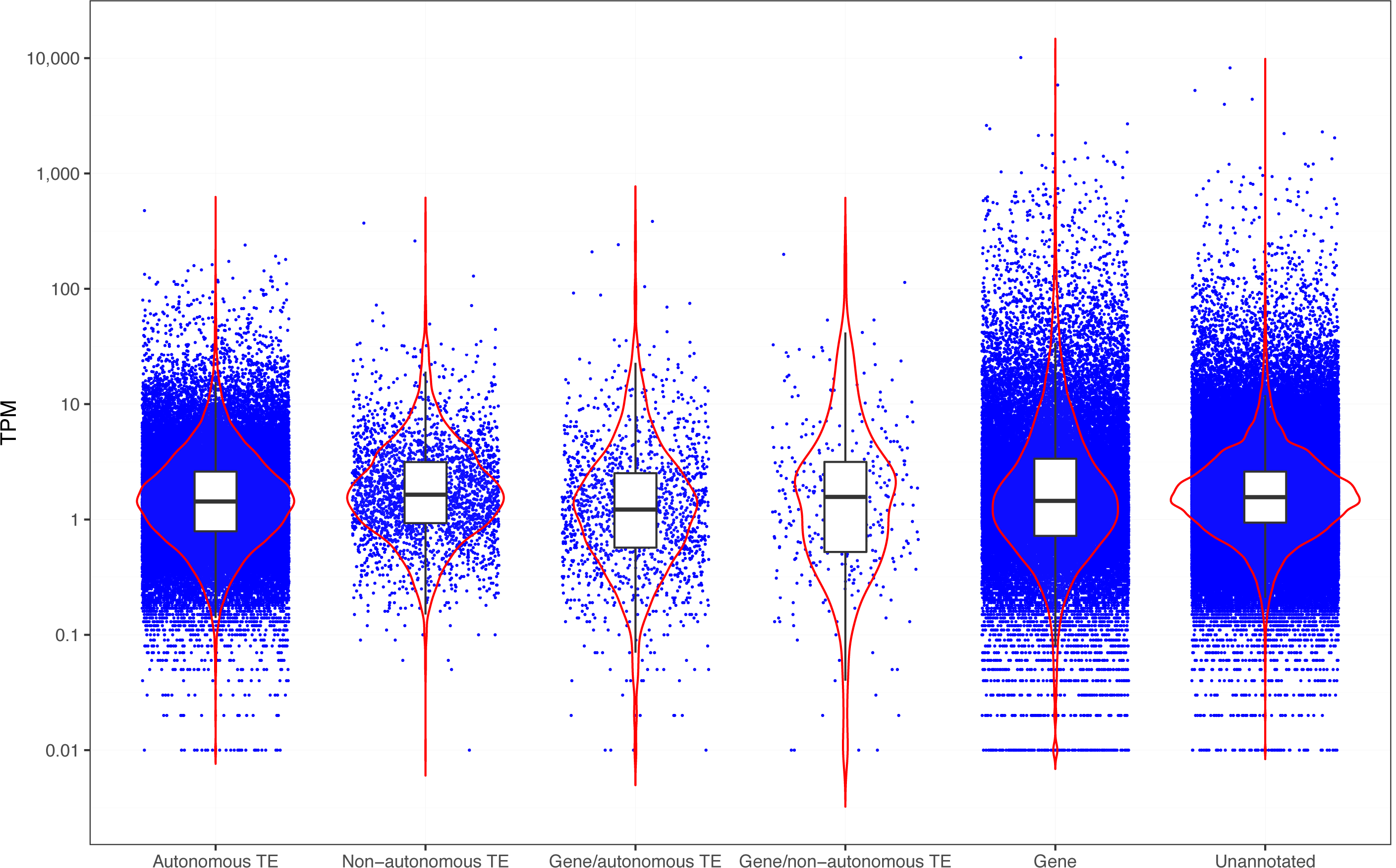
Expression levels of genes and TEs. Black lines and white boxes show median and interquartile range values. Red lines show probability densities. TPM: transcripts per million. TE: transposable element.

Class I TEs (retrotransposons) are over ten times more abundant in the transcriptome than Class II TEs (summed TPM = 130,076 and 10,202, respectively). Within the retrotransposons, DIRS are the most highly expressed, followed by Jockey and L1; these three superfamilies are also the most abundant in the genome. For almost all retrotransposon superfamilies, hundreds to thousands of transcriptome contigs that were <80% identical in sequence to one another were assembled (range 1 – 12,652), suggesting the simultaneous activity of many families within all of the superfamilies in the caecilian somatic transcriptome. Large differences (up to ∼10,000-fold) in expression were detected among the different contigs within superfamilies, suggesting variable expression levels across loci and among families; we interpret this pattern with caution because of the challenges of uniquely mapping short reads to contigs of similar sequence. Within the DNA transposons, Tc1-Mariner, CACTA, and hAT are the most highly expressed superfamilies, and MITEs (transposon derivatives) are expressed at similar levels to these superfamilies, although they lack their own promoters. These four types of sequences are also the four most abundant types of DNA transposons in the genome, although their genomic abundance is not perfectly correlated with their relative expression levels. For the DNA transposons, tens to hundreds of contigs that were <80% identical in sequence to one another were assembled (range 2-421), and up to ∼1000-fold differences in expression were detected among contigs. Overall, a strong correlation was detected between genomic abundance of a TE superfamily and its overall somatic expression level (ρ = 0.879, p < 0.001) (**Figure 4**). Although germline expression data is required to analyze the relationship between TE transcription and TE-activity-driven genome evolution, the somatic data nevertheless provides valuable information on the cellular resources allocated to transcription of a greatly expanded TE community.

**Figure 4.**
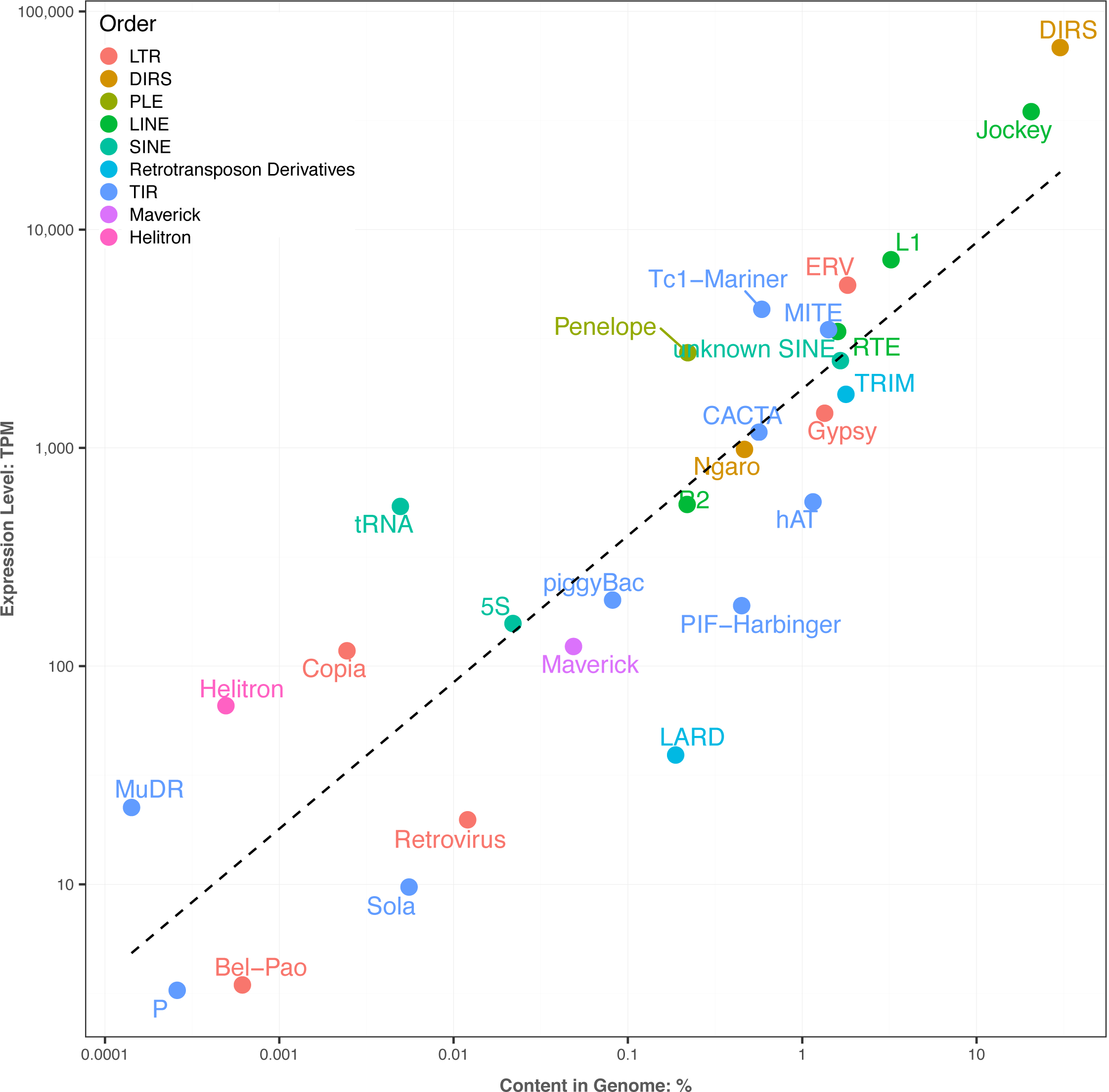
Genomic abundance and somatic expression level of TE superfamilies are strongly correlated (ρ = 0.879, p < 0.001) TPM: transcripts per million.

## Discussion

### Repeat element landscape characterization in large genomes

Large, repetitive genomes have proven difficult to assemble and annotate with the computational power and analytical tools successfully applied to archaeal, bacterial, and smaller eukaryotic genomes [60, 61]. Recent successful genome sequencing efforts aimed at the 32 Gb genome of *Ambystoma mexicanum*, a laboratory model salamander species, leveraged multiple types of data (i.e. optical mapping, short- and long-read genomic sequence data, transcriptomic data, linkage mapping, fluorescence in situ hybridization) and a new assembler designed to minimize compute time and storage requirements [30, 62]. These projects yielded fundamental insights into the structure and evolution of vertebrate chromosomes. They also advanced understanding of the transposons that make up large genomes, adding to research on the 20-Gb Norway spruce and the 22-Gb loblolly and 31-Gb sugar pine [63-65]. This depth of analysis, however, remains infeasible for non-model organisms with large genomes, whose study is nevertheless required for a comprehensive picture of the complex ways in which TEs drive genome biology. Our work affirms the power of low-coverage sequence data to reveal the overall repeat element landscape of large genomes, an approach applied most often in plants (which include the majority of huge genomes) [66, 67]. We argue that this overall landscape, although it lacks the positional information about individual TE insertions that genome assemblies provide, nevertheless contains much information that can be leveraged to identify the evolutionary processes that drive assembly and stability of TE communities.

Repeat element landscapes are informative because they include data on the abundance, diversity, and activity of TEs that make up the overall TE community in a genome. Models of TE dynamics — both formal and informal — predict different values for TE abundance, diversity, and activity depending on levels of purifying selection, silencing, and deletion of TEs. Despite much progress, these forces remain challenging to measure directly. Thus, empirical estimates of TE landscapes provide a feasible alternative by which to validate these models and advance our understanding of TE dynamics in natural systems.

### Repeat element landscapes from large genomes provide tests of models of TE dynamics

Large genomes are especially powerful data points because they represent extreme values of TE abundance, and models of TE dynamics make specific predictions about the effects of TE abundance on TE diversity and activity. We first summarize and highlight the differences among several of these models here (**Figure 5**):

**Figure 5.**
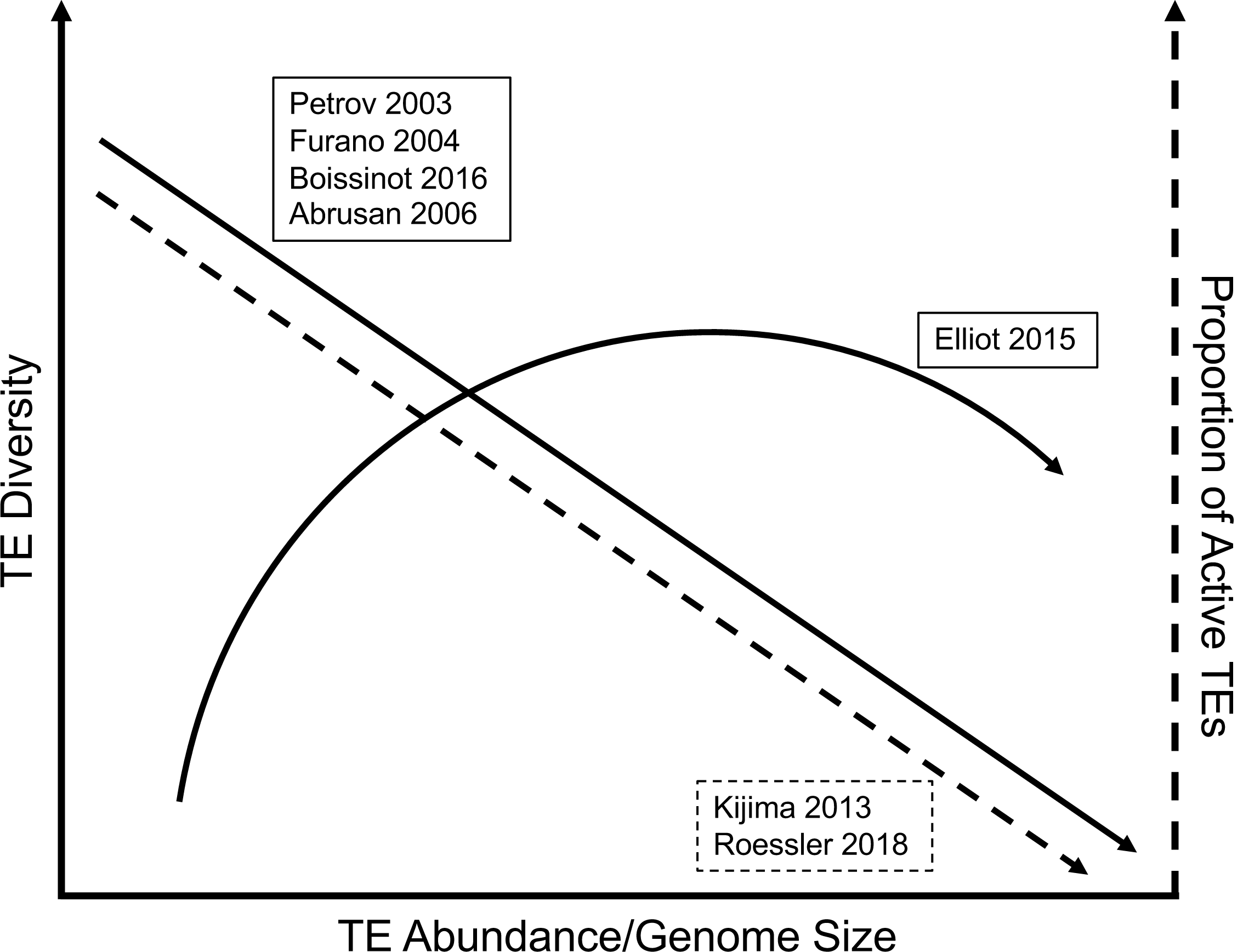
Predicted relationships between TE abundance (genome size), TE diversity, and proportion of active TEs from seven different models

1. Petrov 2003 — TE deletion is caused by ectopic recombination between similar TE sequences. Rates of ectopic recombination/deletion are typically higher in smaller genomes and lower in larger genomes. Thus, smaller genomes are predicted to select for more diverse TE communities, and larger genomes should allow less diverse TE communities [56, 68]. This model predicts an inverse relationship between genome size and TE diversity.
2. Furano 2004 — Because ectopic recombination can cause harmful deletions, it is one of the primary reasons for TEs’ deleterious effects on host fitness. Thus, genomes with lower ectopic recombination/deletion rates are more permissive to TE activity, allowing the accumulation of more TEs (increased genome size) as well as increased TE activity and out-competition of many TE lineages by the lineage that most successfully exploits host replication factors [69]. Like Petrov 2003, this model predicts an inverse relationship between genome size and TE diversity, but for different reasons.
3. Boissinot 2016 — Genomes with lower ectopic recombination/deletion rates have higher levels of insertion of active TE copies into the genome. In addition to yielding a larger genome, this higher number of active TE copies triggers an arms race to control transposition, and the arms race leads to a decrease in diversity (i.e. only one family active at a time) [70]. Like Petrov 2003 and Furano 2004, this model also predicts an inverse relationship between genome size and TE diversity, but for still different reasons.
4. Abrusan 2006 — TE diversity and activity levels were modeled with a system of differential equations that includes parameters for the number of TE strains, the number of active TE insertions, TE replication rates, the strength of specific silencing of TEs (representing small-RNA-mediated silencing), cross-reactivity of silencing, and TE inactivation by mutation or selection [71]. Although their model did not specifically address genome size, it did predict that increased genome size would be associated with decreased TE diversity if a) larger genomes harbor more active TE copies, and b) cross-reactive silencing exists among TEs. Under these conditions, competition among the TEs to evade cross-reactive silencing would lead to decreased TE diversity. Cross-reactive silencing in this model is not sequence-specific; this is relevant because silencing of TEs by small-RNA-mediated silencing (e.g. the piRNA pathway) is sequence-specific, but can have some off-target effects. These off-target effects are predicted to have the opposite effect on TE diversity than non-sequence-specific cross-reactive silencing; they should select for higher TE diversity. Overall, the predictions for genome size and TE diversity from this model are complex, depending on the relative strengths of specific TE silencing, off-target specific TE silencing, and cross-reactive (i.e. sequence-independent) silencing.
5. Elliot 2015 — Based on empirical comparisons across genomes of different sizes, TE diversity was proposed to increase with TE abundance until genomes reach moderate size, but extremely large genome sizes were proposed to reflect the proliferation of only a subset of TE diversity by unspecified mechanisms [72]. This predicts an inverse relationship between genome size and TE diversity at the largest genome sizes.
6. Kijima 2013 — TE evolution was modeled using a population genetic simulation framework that includes parameters for transposition, TE deletion, purifying selection on TE copy number (genome size), and degeneration into inactive copies [73]. When copy number selection is strong (i.e. genome size remains small), the total number of TEs is lower, but the proportion of active copies of TEs is higher. When copy number selection is weak (i.e. genome size is allowed to increase), the total number of TEs is higher, but the proportion of active copies of TEs is lower. This reflects competition among TEs to occupy limited available spaces in the genome. This model does not consider TE diversity — it models only a single TE strain — but it predicts an inverse relationship between genome size and proportion of the total TE community that is actively transposing. Interestingly, they find that excision (deletion) rate is not a predictor of copy number.
7. Roessler 2018 — TE evolution was modeled using ordinary differential equations including parameters for TE transposition, RNA-mediated TE silencing, TE deletion, and TE copy number (genome size) [74]. This model predicts that, under low rates of TE deletion, TE copy number and genome size increase, and the proportion of active TEs goes down because the host organism can use the accumulating TE sequences as templates for producing more small silencing RNAs and, thus, inactivate a higher proportion of TE sequences. Like Kijima 2013, this model predicts an inverse relationship between genome size and proportion of the total TE community that is actively transposing, but for different (albeit complementary) reasons.

Does the TE landscape of the large caecilian genome — with its high levels of TE abundance and low levels of TE ectopic recombination/deletion — fit the predictions of these models or allow discrimination among them? Most share a prediction of decreased TE diversity in large genomes. Measured at the coarse-grained level of number of superfamilies present (i.e. taking into account richness only) [72], *I. bannanicus* does not fit this prediction; at least 25 TE superfamilies are present in the genome (as detected by our genomic and transcriptomic analysis). However, genome expansion in *I. bannanicus* is correlated with high DIRS/DIRS and LINE/Jockey superfamily abundance, consistent with Elliot 2015’s prediction that gigantic genomes would reflect proliferation of a limited subset of all TEs. This expansion decreases evenness, despite the maintenance of high richness; this is exactly the type of change in overall diversity that is captured by the indices we advocate here.

Comparing the diversity indices calculated for *I. bannanicus* with the 11 other vertebrate genomes (Table 3) allows a direct test of the relationship between genome size (i.e. TE abundance) and TE diversity. Because the genomes included were analyzed with different sequencing depths, we favor the Gini-Simpson Index as it is less affected by rare species (TE superfamilies), which are more likely missed in the low-coverage datasets (e.g. *I, Zisupton, Kolobok, Academ*, and *Crypton*; Table 2). Consistent with model predictions, the smallest genome (*T. rubripes*) has the highest TE diversity, and the three most diverse genomes (*T. rubripes, A. carolinensis*, and *X. tropicalis*) are three of the four smallest (Table 3). However, among the large amphibian genomes — *I. bannanicus* and the five salamanders — there is no relationship between TE abundance and diversity. Furthermore, the chicken genome is the least diverse, and it is the second-smallest.

However, the lack of relationship between TE abundance and diversity, measured here at the TE superfamily level for 11 species, does not necessarily refute the models of TE dynamics that predict decreased TE diversity with increased TE abundance. Diversity exists within TE superfamilies as well; TE families are typically operationally defined based on Wicker’s 80/80/80 rule, and subfamilies can be further split based on pairs of substitutions overrepresented in TE alignments that are unlikely to have arisen independently by chance [7, 75]. It is not yet clear what levels of sequence divergence translate into functionally relevant “TE diversity” in the models summarized above. More specifically, TE diversity implies: 1) TE sequences that have diverged beyond the ability to ectopically recombine in Petrov 2003, 2) TE sequences that have diverged enough to differ in ability to monopolize host replicative resources in Furano 2004, 3) TE sequences that have diverged enough to (sequentially) out-evolve host silencing machinery in Boissinot 2016, and 4) TE sequences that have diverged enough to differ in their silencing by cross-reactive (i.e. non-sequence-specific) or off-target (i.e. sequence-specific, but tolerant of mismatches) TE silencing mechanisms in Abrusan 2006. We still lack sufficient information about TE silencing to define the levels of sequence divergence likely to accompany these changes in TE dynamics. Thus, it is not yet clear whether diversity indices are best focused at the TE superfamily, family, or subfamily levels. As an example, the chicken genome is the least diverse measured here at the level of TE superfamilies because CR1 elements dominate the genome; however, diversity exists within the CR1 elements that may be functionally relevant [76]. To move the field forward, we advocate using Shannon and Simpson indices at the levels of TE family and subfamily (in addition to superfamily) when datasets allow. When this is impossible — for example, when working with low-coverage shotgun data from gigantic genomes like *I. bannanicus* — we advocate calculating diversity indices at the superfamily level, but also reporting the numbers of genomic and transcriptomic contigs < 80% identical as a tractable within-superfamily approximation of TE diversity (Table 2). This measure is analogous to species richness and lacks information on evenness (because of the challenges of uniquely mapping short reads to contigs of similar sequence), so it is less informative than diversity indices. However, the reporting of this measure by researchers studying diverse organisms would allow progress towards rigorously testing the relationship between genome size and TE diversity. Furthermore, it may identify specific taxa as appropriate models to examine evolutionary changes in TE silencing pathways. For example, *I. bannanicus* has a large genome but appears to maintain a high number of TE families (Table 2), suggesting that its TE silencing machinery includes high levels of off-target silencing [71].

In addition to predicting low TE diversity, models of TE dynamics predict a decreased proportion of active TEs as TE abundance and genome size increase. Of the 19 caecilian TE superfamilies for which amplification histories were examined, 17 appear to have ongoing activity (Figure 1). These results are largely corroborated by the (albeit somatic) expression data, although SINE/7SL and LINE/R2 show conflicting patterns in the genomic and transcriptomic data (Figure 1, Table 2). TE expression is necessary, but insufficient, for TE activity, but it is a tractable proxy for TE activity. Taken together, these datasets suggest near-complete activity at the TE superfamily level in the *I. bannanicus* genome. At the levels of TE family or individual insertions, activity is difficult to assess with our data; however, the presence of multiple transcriptome contigs <80% identical within superfamilies minimally suggests the expression of multiple families. Our recommendation that researchers report the number of transcriptomic TE contigs < 80% identical will also allow progress towards rigorously testing the relationship between genome size and TE activity, as will adoption of recent methods to measure locus-specific expression when datasets allow [77].

Overall, ∼15% of all somatic tissue transcripts of *I. bannanicus* are TEs (Table 4). Comparing overall levels of TE expression across different genome sizes remains difficult because TE expression in general is understudied [77], transcriptome size differences that accompany genome size differences are typically not quantified [78], and TE annotation and expression quantification methods vary across studies [38, 79-81]. As another step towards testing the relationship between genome size and TE activity, we advocate annotation of both autonomous and non-autonomous TE transcripts and reporting of expression levels of TEs and endogenous genes (Table 2, Table 4, Figure 3).

Taken together, our work lays a foundation for comparative genomic analyses that link properties of TE communities — abundance, diversity, and activity — to genome size evolution. Such analyses, in turn, will reveal whether the divergent TE assemblages found across convergent examples of genomic gigantism reflect more fundamental shared features of TE/host genome evolutionary dynamics.

## Materials and Methods

### Specimen information

We collected a single male adult caecilian (*I. bannanicus*) from the species’ type locality (E101.3887, N21.8724) in Mengxing County, Yunnan province, China. The individual had a total body length of 16.0 cm and a body mass of 4.8 g. Following dissection, the carcass was fixed in formalin and transferred to 70% ethanol.

### Genome size estimation

Blood smears were prepared from the formalin-fixed *I. bannanicus* specimen as well as a formalin-fixed salamander (*Plethodon cinereus*) with an appropriate genome size to serve as the reference standard (22.14 Gb) [82]. Blood cells were pipetted onto glass microscope slides and air-dried, then hydrated for three minutes in distilled water. Slides were 1) hydrolyzed in 5N HCl for 20 minutes at 20°C and washed three times in distilled water for one minute each, 2) stained with Schiff’s reagent in a Coplin jar for 90 minutes at 20°C, 3) soaked in three changes of 0.5% sodium metabisulfite solution for five minutes each and rinsed in three changes of distilled water for one minute each, and 4) dehydrated in 70%, 95%, and 100% ethanol for one minute each, air-dried, and mounted in immersion oil and cover glass.

The stained slides were photographed using an Olympus BX51 compound microscope fitted with a Spot Insight 4 digital camera for image analysis. Stained nuclei were photographed under 100x oil immersion and the integrated optical densities were measured using ImagePro software. Genome size for *I. bannanicus* was calculated by comparing the mean optical density to that of the reference standard, *P. cinereus*.

### Genomic shotgun library creation, sequencing, and assembly

Total DNA was extracted from muscle tissue using the modified low-salt CTAB extraction of high-quality DNA procedure [83]. DNA quality and concentration were assessed using agarose gel electrophoresis, a NanoDrop Spectrophotometer (Thermo Scientific), and a Qubit 2.0 Fluorometer (ThermoFisher). A PCR-free library was prepared using NEBNext Ultra DNA Library Prep Kit for Illumina. Sequencing was performed on two lanes of a Hiseq2500 platform (PE250). Library preparation and sequencing were performed by the Beijing Novogene Bioinformatics Technology Co. Ltd. Raw reads were quality-filtered and trimmed of adaptors using Trimmomatic-0.36 [84] with default parameters. In total, the genomic shotgun dataset included 7,785,846 reads. After filtering and trimming, 7,275,133 reads covering a total length of 1,635,569,256 bp remained. Thus, the sequencing coverage is 0.134. Filtered, trimmed reads were assembled into contigs using dipSPAdes 3.11.1 [85] with default parameters, yielding 130,417 contigs with an N50 of 740 bp and a total length of 1,560,938,851 bp.

### Mining and classification of repeat elements

The PiRATE pipeline was used as in the original publication [44], including the following steps: 1) Contigs representing repetitive sequences were identified from the assembly using similarity-based, structure-based, and repetitiveness-based approaches applied non-sequentially. The similarity-based detection programs included RepeatMasker [86] and TE-HMMER [87]. The structure-based detection programs included MITE-Hunter [88], SINE-finder [89], HelSearch [90], LTRharvest [91], and MGEScan-nonLTR [92]. The repetitiveness-based detection programs included TEdenovo [93] and RepeatScout [94]. 2) Contigs representing repeat family consensus sequences were also identified from the cleaned, filtered, unassembled reads with dnaPipeTE [95], which uses Trinity on subsamples of single-end reads to produce sets of related repeat consensus sequences (e.g. representing multiple subfamilies within a TE family). 3) Contigs identified by each individual program in Steps 1 and 2, above, were filtered to remove those <100 bp in length and clustered with CD-HIT-est [96] to reduce redundancy (100% sequence identity cutoff). This yielded a total of 62,699 contigs. 4) All 62,699 contigs were then clustered together with CD-HIT-est (100% sequence identity cutoff), retaining the longest contig and recording the program that classified it. 1860 contigs were filtered out at this step, and the majority (1669) were contigs identified by RepeatMasker and TE-HMMER that were identical in sequence but differed in length. 5) Repeat contigs were annotated as TEs to the levels of order and superfamily in Wicker’s hierarchical classification system [7], modified to include several recently discovered TE superfamilies, using PASTEC [45] and were checked manually to filter chimeric contigs and those annotated with conflicting evidence. 6) All classified repeats (“known TEs” hereafter), along with the unclassified repeats (“unknown repeats” hereafter) and putative multi-copy host genes, were combined to produce a caecilian-derived repeat library.

### Characterization of the overall repeat element landscape

Overlapping paired-end reads were merged using PEAR v.0.9.11 [97] with the following parameter values based on our library insert size and trimming parameters: min-assemble-length 36, max-assemble-length 490, min-overlap size 10. After merging the remaining paired-end reads, 6,628,808 shotgun reads remained, with an average and a total length of 236 and 1,560,938,851 bp, respectively. To calculate the percentage of the caecilian genome composed of different TEs, the shotgun reads (including both merged reads and singletons) were masked with RepeatMasker v-4.0.7 using two versions of our caecilian-derived repeat library: one that included the unknown repeats and one that excluded them. In both cases, simple repeats were identified using the Tandem Repeat Finder module implemented in RepeatMasker. The overall results were summarized at the levels of TE class, order, and superfamily. For each superfamily, we then collapsed the contigs to 95% and 80% sequence similarity using CD-HIT-est to provide an overall view of within-superfamily diversity; 80% is the sequence similarity threshold used to define TE families [7].

### TE community diversity

Diversity of the overall transposable element community in *I. bannanicus* was summarized using the Shannon index H′ = − ∑P_*i*_ ln(P_*i*_)and the Simpson index 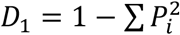 (i.e. the Gini-Simpson index), where P_*i*_ is the proportion of sequences belonging to TE superfamily *i* [51, 52]. In analogous applications of these diversity indices to ecological communities, P_*i*_ is the proportion of individuals that belong to species P_*i*_. To provide context for the *I. bannanicus* results, Shannon and Simpson indices were also calculated for other vertebrate genomes representing diversity in genome size as well as type of dataset. *Takifugu rubripes* (pufferfish, 0.4 Gb), *Gallus gallus* (chicken, 1.3 Gb), *Xenopus tropicalis* (Western clawed frog, 1.7 Gb), *Anolis carolinensis* (green anole lizard, 2.2 Gb), and *Homo sapiens* (human, 3.1 Gb) all have full genome assemblies. For these five species, the perl script parseRM.pl [98] was used to parse the raw output files downloaded from www.repeatmasker.org and obtain the percentage of the genome occupied by each identified superfamily; ambiguous classifications (i.e. to the level of order or class) were excluded. *Ambystoma mexicanum* (the axolotl, a model salamander, 32 Gb), which has a much larger genome and, consequently, less complete genome assembly, was also included; percentages of the genome occupied by each identified superfamily were obtained from a previous study [30]. Finally, four other salamanders that encompass a range of genome sizes were included, each represented by low-coverage genome-skimming shotgun data: *Desmognathus ochrophaeus* (15 Gb), *Batrachoseps nigriventris* (25 Gb), *Aneides flavipunctatus* (44 Gb), and *Cryptobranchus alleganiensis* (55 Gb). Percentages of each genome occupied by identified superfamilies were obtained from a previous study [32].

### Amplification history of transposable element superfamilies

To summarize the overall amplification history of TE superfamilies and test for ongoing activity, the perl script parseRM.pl [98] was used to parse the raw output files from RepeatMasker (.align) and report the sequence divergence between each read and its respective consensus sequence (parameter values = -l 50,1 and -a 5). The repeat library used to mask the reads comprised the 50,471 TE contigs classified by the PiRATE pipeline and clustered at 100% sequence similarity. Each TE superfamily is therefore represented by multiple consensus contigs that represent ancestral sequences likely corresponding to the family and subfamily TE taxonomic levels (i.e. not the distant common ancestor of the entire superfamily). For each superfamily, histograms were plotted to summarize the percent divergence of all reads from their closest (i.e. least divergent) consensus sequence. These histograms do not allow the delineation between different amplification dynamics scenarios (i.e. a single family with continuous activity versus multiple families with successive bursts of activity). Rather, these global overviews were examined for overall shapes consistent with ongoing activity (i.e. the presence of TE loci <1% diverged from the ancestral sequence and a unimodal right-skewed, J-shaped, or monotonically decreasing distribution).

### Ectopic recombination-mediated deletion of Gypsy and DIRS elements

All genomic contigs > 3000 bp in length that were annotated to LTR/Gypsy were *de novo* annotated using LTRpred to identify terminal and internal sequences [99]. Internal and terminal sequences were further confirmed by manually checking for internal TE domains using NCBI BLASTx (https://blast.ncbi.nlm.nih.gov/Blast.cgi?PROGRAM=blastx&PAGE_TYPE=BlastSearch&LINK_LOC=blasthome) and for LTR sequences using the NCBI-Blast2suite to align each contig sequence against itself. DIRS/DIRS superfamily elements have a different structure than other LTR retrotransposons; their terminal repeats are inverted. However, because they also include internal sequences complementary to the terminals that facilitate rolling-circle amplification [54, 55], their structure includes direct repeats that are expected to undergo ectopic recombination to eliminate much of the internal sequence and one copy of the direct repeat sequence, although to our knowledge this has not been previously investigated. Although these deletions would not produce canonical solo LTRs, they, too, would produce elevated abundances of terminal sequences relative to internal sequences. Typical DIRS structure was confirmed visually and by using the NCBI-Blast2suite to align each contig sequence against itself, and contigs that lacked the complete structure were removed from further analysis. Internal sequences for both superfamilies were conservatively defined to be bounded by the first and last TE domains. This yielded a total of nine DIRS/DIRS contigs and 17 LTR/Gypsy contigs. DIRS/DIRS contigs had an average terminal sequence length of 150 bp (range 61-343) and an average internal sequence length of 5586 bp (range 4810-6012). LTR/Gypsy contigs had an average terminal sequence length of 744 bp (range 127 – 3267) and an average internal sequence length of 1976 (range 243 – 4306). To estimate levels of terminal sequences (LTRs or TIRs) relative to internal sequences, genomic shotgun reads were mapped to the whole genome assembly using bowtie2 in local alignment mode with very-sensitive-local preset options and otherwise default parameters, increasing the G-value from the default of 20 to 30, 40, and 50 to increase minimum alignment length for reads [100]. This analysis was performed twice: once treating all reads as unpaired and once using merged paired-end reads plus unmerged reads. Average read depths across the terminal and internal portion in each of the 26 focal DIRS/DIRS and LTR/Gypsy contigs were estimated by scaling the number of hits by the lengths of the terminal and internal region. From these estimates, the total terminal-to-internal sequence ratio (TT:I) was calculated for each contig. In the absence of ectopic recombination mediated by terminal repeats, this ratio would be 1:1; increasing levels of ectopic recombination would produce ratios > 1:1. We compared the results obtained for the caecilian with similar analyses that included gigantic salamander genomes as well as vertebrates with more typical (i.e. smaller) genomes [33].

### Transcriptome library creation, sequencing, assembly, and TE annotation

Total RNA was extracted separately from heart, brain, liver, and tail tissue using TRIzol (Invitrogen). For each sample, RNA quality and concentration were assessed using agarose gel electrophoresis, a NanoPhotometer spectrophotometer (Implen, CA, USA), a Qubit 2.0 Fluorometer (ThermoFisher), and an Agilent BioAnalyzer 2100 system (Agilent Technologies, CA, USA) requiring an RNA integrity number (RIN) of eight or higher. Equal quantities of RNA from these four tissues were pooled to build a single transcriptome library. Sequencing libraries were generated using the NEBNext Ultra RNA Library Prep Kit for Illumina following the manufacturer’s protocol. After cluster generation of the index-coded samples, the library was sequenced on one lane of an Illumina Hiseq 4000 platform (PE 150). Library preparation and sequencing were performed by the Beijing Novogene Bioinformatics Technology Co. Ltd. Transcriptome sequences were filtered using Trimmomatic-0.36 (Bolger, et al. 2014) with default parameters. Remaining reads were assembled using Trinity 2.5.1 (Grabherr, et al. 2011). In total, 34,980,300 transcriptome reads were obtained, with a total length of 5,247,045,000 bp. After filtering, 34,417,105 reads remained, with a total length of 5,027,542,505 bp. The assembly produced 348,822 contigs (i.e. putative assembled transcripts) with the min, N50, max, and total length of contigs equal to 201; 357; 32,175; and 249,943,402 bp, respectively. Of these, 289,380 had expression levels of TPM ≥ 0.01 and were analyzed further.

To annotate transcriptome contigs containing autonomous TEs, BLASTx was used against the Transposable Element Protein Database (RepeatPeps.lib, downloaded from https://github.com/rmhubley/RepeatMasker/blob/master/Libraries/ on April 20, 2019) with an e-value cutoff of 1e-10. To annotate contigs containing non-autonomous TEs, RepeatMasker was used with our caecilian-derived genomic repeat library of non-autonomous TEs (LARD-, TRIM-, MITE-, and SINE-annotated contigs; Table 2) and the requirement that the transcriptome/genome contig overlap was >80 bp long, >80% similar in sequence, and covered >80% of the length of the genomic contig. Contigs annotated as conflicting autonomous and non-autonomous TEs were filtered out. To yield a rough estimate of the number of active TE families per superfamily, CD-HIT-est was used to cluster the contigs annotated to each superfamily at the level of 80% sequence similarity.

To identify contigs that contained an endogenous caecilian gene, the Trinotate annotation suite was used with e-value cutoffs of 1e-10 and 1e-5 for BLASTx and BLASTp against the SwissProt database, respectively, and 1e-5 for HMMER against the Pfam database [57]. To identify contigs that contained both a TE and an endogenous caecilian gene (i.e. putative cases where a TE and a gene were co-transcribed on a single transcript), all contigs that were annotated both by RepeatPeps and Trinotate were examined, and the ones annotated by Trinotate to contain a TE-encoded protein (i.e. the contigs where RepeatPeps and Trinotate annotations were in agreement) were not further considered. The remaining contigs annotated by Trinotate to contain a non-TE gene (i.e. an endogenous caecilian gene) and also annotated either by RepeatPeps to include a TE-encoded protein or by RepeatMasker to include a non-autonomous TE were identified for further examination and expression-based analysis.

### Transposable element expression

To generate a point estimate of overall TE expression in the somatic transcriptome, transcript abundance levels were quantified with RSEM (because of its capacity to model multi-mapping reads) using the Bowtie short-read aligner. Transcriptome contigs with TPM < 0.01 were filtered out. To yield TE-superfamily-wide expression level estimates, TPM values were summed across all contigs annotated to the same TE superfamily. For comparison, TPM values were summed for all endogenous (i.e. non-TE) caecilian genes. Pearson’s correlation coefficient was used to test for a relationship between genomic TE abundance (measured as log-transformed percentage of the genome occupied per TE superfamily) and TE expression level (measured as log-transformed total TPM per TE superfamily). We note that with only a single sample, any more detailed analyses of expression levels are not appropriate. Contigs annotated to contain both TEs and endogenous caecilian genes were excluded from these analyses. Instead, these putative TE/gene contigs were ranked by expression level, and the 20 most highly expressed were examined by eye to determine the spatial relationship between the TE and gene BLAST results producing the annotations. Nine contigs with apparently spurious TE annotations (seven of which reflected a single likely mis-annotation of an LTR/Pao protein in the RepeatPeps database) were reclassified as endogenous genes, and the remaining contigs were characterized as having the TE 1) on the same or different strand as the gene, and 2) upstream or downstream of the gene. Finally, TPM values were summed across all putative TE/gene contigs to yield a global estimate of expression levels of TE/gene combinations that are co-transcribed on a single transcript.

## Ethical Statement

The study specimen was collected and dissected following Animal Care & Use Protocols approved by Chengdu Institute of Biology, Chinese Academy of Sciences.

## Data availability

Genomic shotgun and transcriptome sequences have been deposited in the National Genomics Data Center (accession numbers: SAMC207357, SAMC207358).

## Authors’ contributions

JW and RM designed and supervised the experiments. JW, MI, HW, and CS participated in bioinformatic analysis. MI and SS participated in genome size measurement. YG collected the sample from the field. JJ and JL advised on data analysis. JW and RM wrote the manuscript with input from all authors. All authors read and approved the final manuscript.

## Acknowledgments

This work was supported by the National Natural Science Foundation of China (31570391 to W.J.), the National Key Programme of Research and Development (Ministry of Science and Technology, 2016YFC0503200 to J.J.P). the Western Light Foundation of the Chinese Academy of Sciences (Y3C3011100 to W.J.), the National Science Foundation (1911585 to R.L.M.). We gratefully acknowledge Xungang Wang, Liming Chang, Aurélie Kapusta, Lusha Liu, Yiwei Zeng, and Shan Xiong for target tissue dissection; Daniel B. Sloan, Yufeng Wang, Zhiqiang Wu, Evan Forsythe, and Ava Lambert for their expertise in facilitating data analysis; and Jérémy Berthelier for assistance running PiRATE.

